# A two-signal, two-receiver signalling system: Snake mobbing in the Arabian babbler

**DOI:** 10.64898/2026.08.25.747098

**Authors:** Marie Guggenberger, Oded Keynan, Yossi Yovel

## Abstract

Mobbing is a collective antipredator behaviour in which animals approach and harass a threat while coordinating signals to alert conspecifics, recruit allies, or deter predators. In group-living Arabian babblers (*Argya squamiceps*), snake mobbing relies on two signals that differ in receivers: acoustic “zwick” calls inaudible to the snake and visual wing-lift displays perceived by both conspecifics and the snake, raising the question of whether each signal has one receiver or involves a more complex structure. We elicited snake-mobbing events with dummy vipers and recorded babbler groups using a synchronized acoustic camera enabling individual caller identification and postural analysis. We examined mob formation, the effect of group size on signalling, and the interplay between the two signal types. First signalling (recruitment) order was independent of sex, age and rank. Individual call rate decreased as group size increased, supporting the hypotheses of social buffering and predation risk dilution. Mobbing individuals responded faster (with signalling) to a joint visual-acoustic conspecific display and an onset of a visual display than to an acoustic-only signal. When coupled, wing-lifts significantly altered vocal characteristics, decreasing peak frequency and increasing call rate, thus strengthening the acoustic signal. Moreover, calls emitted toward the end of the wing-lift display exhibited a stronger drop in peak frequency. Together, these findings demonstrate the multi-functional role of multimodal mobbing signals during a risky cooperative task balancing social communication and predator deterrence.

## Introduction

Communication plays a central role in many forms of collective behaviour, allowing individuals to coordinate their actions and respond adaptively to dynamic environmental conditions (Bradbury and Vehrencamp, 1998; Higham and Hebets, 2013; Liao et al., 2024; Sebeok, 1965). Signals can transmit information about resources, threats, or the actions of other group members, enabling animals to synchronize movements, recruit partners, and adjust their behaviour based on the state of the group (Bradbury and Vehrencamp, 1998; Hölldobler, 1999). Social insect and vertebrate communication systems often integrate multiple sensory channels, such as acoustic, visual, or chemical signals, which together allow information to spread efficiently through groups and facilitate coordinated responses (Partan and Marler, 2005; Sebeok, 1965).

Group behaviour is often shaped by social phenomena such as social contagion and social buffering (Hennessy et al., 2009; Oliveira and Faustino, 2017). Individuals within a group may therefore respond to conspecific behaviour by either copying the signal or by reducing their own signalling effort as others join. Through these mechanisms, communication enables individuals to transform local information into coordinated group-level behaviour, a process that is particularly evident in antipredator contexts, where signalling can rapidly recruit allies and organize defensive responses, of which mobbing represents a striking example (Flasskamp, 1994).

Mobbing is a collective antipredator behaviour with immediate survival consequences, in which animals approach a threat while often simultaneously coordinating vocalizations and movements (Abolins-Abols and Ketterson, 2017; Chandler and Rose, 1988; Courter and Ritchison, 2012; Ostreiher, 2003). The behaviour generally involves prey grouping to deter predators, assess risk, and alert others to danger (Carlson and Slabbekoorn, 2026; Flasskamp, 1994; Graw and Manser, 2007). Beyond immediate survival, it likely serves secondary purposes such as signalling physical strength and social rank and teaching young individuals about risk assessment (Curio et al., 1978; Graw and Manser, 2007; Lima and Dill, 1990; Maklakov, 2002).

Variation in individual traits such as age-sex category or social rank can systematically bias risk-taking behaviour, for example, the decision of whether and when to join a mob, influencing both individual decision-making and the dynamics of collective responses (Blaszczyk, 2017; Brand et al., 2023; Francis et al., 1989; Graw and Manser, 2007; Hogstad, 1988; Smit and Robbins, 2025). In captive groups of common ravens (*Corvus corax*), dominant individuals produced scolding calls earlier and for longer durations than subordinates (Blum et al., 2022). Those rank differences disappeared in isolation, indicating that social dynamics influence antipredator signalling effort within groups (Blum et al., 2022). In collective contexts, social information use further modulates individual contributions: social buffering, demonstrated in zebrafish, diminishes an individual’s response to a detected threat (e.g., alarm substance) when visual and olfactory cues of relaxed conspecifics signal its absence (Oliveira and Faustino, 2017). Social contagion has the opposite effect: it triggers an alarm response from conspecifics’ distress cues alone, without direct threat detection (Oliveira and Faustino, 2017).

Communication signals are key to successful mobbing. In birds, mobbing often combines acoustic and visual displays: Anna’s hummingbirds (*Calypte anna*) actively circle raptors while producing short, high-pitched notes and directing their bill toward the predator’s eyes (Altmann, 1956), and American robins (*Turdus migratorius*) combine calls with wing-flicking, tail-wagging, and restless hopping in proximity to predators (Shedd, 1978). Such multimodal signalling allows individuals to transmit richer or more robust information than a single modality alone could convey (Gupta et al., 2022; Hölldobler, 1999; Liao et al., 2024; Partan and Marler, 2005; Sebeok, 1965). When channels influence each other, body movements can, for example, shift the acoustic properties of a signal, altering parameters such as frequency, rate, or duration in ways that may carry additional information about threat urgency or caller motivation (Caldart et al., 2022; Hoepfner and Goller, 2013; Longo et al., 2019; Vorperian et al., 2015; Yorzinski and Vehrencamp, 2009; Zahavi, 1982). In human communication, assertive or bullying displays utilize a multichannel system where expansive postures, such as head raising and torso straightening, physically facilitate the production of higher vocal pitch and loudness (Berry et al., 2022).

The exact function of the various signals used during mobbing is hard to decipher and also dependent on the physiology of the receiver. Snakes, for example, unlike most other predators, lack an external ear and therefore cannot perceive acoustic signals above 1000 Hz (Christensen et al., 2012). Calls frequently recruit additional participants, both from within the group and from neighbouring groups, and even sometimes across species (Coye and Dutour, 2025; Dutour et al., 2017; Kern and Radford, 2016). Beyond recruiting conspecifics, calls may also be directed at the predator itself. Under the pursuit-deterrent hypothesis, they signal that the caller has detected it, potentially reducing its motivation to attack (Caro, 1995; Sommer et al., 2012; Woodland D. J. et al., 1980). Visual displays during predator encounters can serve the same dual role, though separating deterrence from recruitment remains difficult. While visual postural displays are typically effective at short range and may target nearby animals primarily, acoustic signals can propagate over longer distances and therefore reach dispersed group members and more distant predators (Bradbury and Vehrencamp, 1998).

Mobbing is difficult to study because it typically involves many individuals converging from different directions, often at high density while moving fast and vocalizing nearly at the same time, making it hard to track individuals and identify vocalizers. This is further complicated under wild conditions, where mobbing cannot easily be elicited artificially. To overcome these challenges, we used a combination of a suitable animal model and a novel recording approach.

Arabian babblers (*Argya squamiceps*) are cooperatively breeding, group-living passerines inhabiting deserts. They actively use a variety of acoustic signals for group coordination and engage in group mobbing against several predator types (Maklakov, 2002; Naguib et al., 1999; Sommer et al., 2012). A typical group consists of one dominant breeding pair and several related or non-related helpers, who assist in feeding the young and defending against predators (Dragić et al., 2021; Ridley and Huyvaert, 2007). When an Arabian babbler discovers a snake, it begins emitting “zwick” calls, which attract other babblers. Individuals then approach and spread around the snake while continuing to call (Guggenberger et al., 2024; Maklakov, 2002). Some birds occasionally adopt a specific display posture — spreading the tail and lifting and fanning the wings — while circling the predator (Zahavi, 1997). Babblers are highly suitable for studying multimodal signalling during mobbing because a mock snake can repeatedly and reliably elicit mobbing (Guggenberger et al., 2024; Maklakov, 2002; Ostreiher, 2003). The fact that they mob on the ground (and not in flight) makes the tracking of individuals more accessible. To elicit mobbing events, we used a dummy viper and recorded babbler groups simultaneously with a synchronized acoustic camera (SoundCam 2.0) and a high-resolution RGB video camera positioned approximately three meters from the model (Fig. 1A; Supplementary Video 1 and 2), enabling precise individual caller identification through sound localization and postural tracking (Fig. 1B) to disentangle individual contributions within a naturally coordinated group response. We used this system to examine babbler mobbing behaviour across five levels, from mob formation to the fine-grained interaction between signals, asking: (1) whether individual traits, age-sex category and social rank, predict the order in which birds actively (showing mobbing behavior) join the mob; (2) how the multimodal signal changes the response latency of others during active mobbing; (3) how individual investment in acoustic and visual signalling scales with group size; (4) what the functional and physiological relationship between “zwick” calls and wing-lift displays is; and (5) how the acoustic structure of calls changes depending on the relative position within a simultaneous wing-lift display, and how these changes evolve within and across successive displays.

**Fig. 1.**
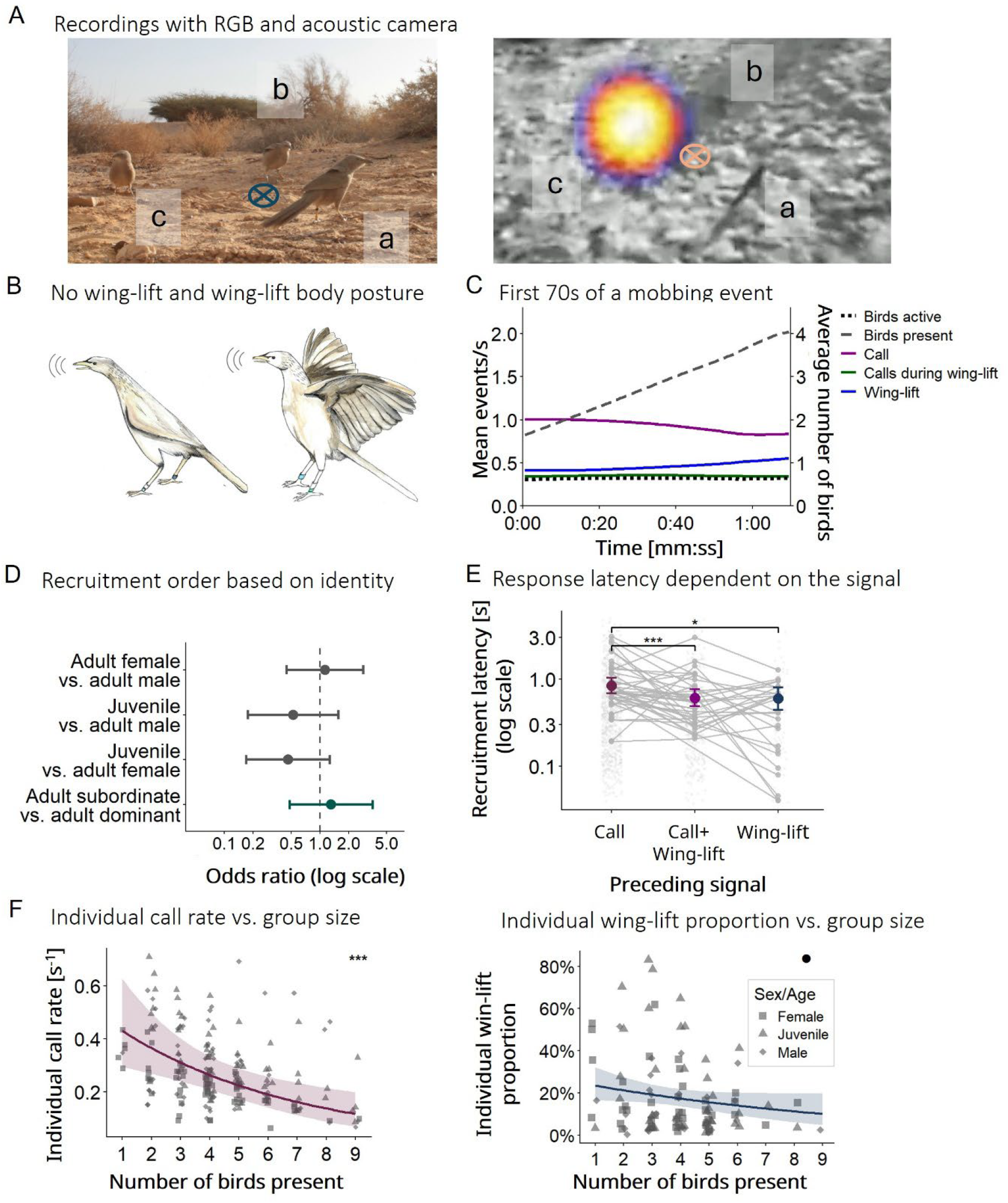
Experimental setting, mobbing dynamics, and recruitment and individual mobbing behaviour. **A, left**, RGB camera recording of babblers with identifiable leg-band colours (individuals a–c are marked). **Right**, zoomed-in acoustic camera frame during a vocalization by bird c, visualized as a sound-pressure gradient from high dB (white) to low dB (blue). **B**, Arabian babbler without (left) and during (right) the wing-lift display. **C**, mean events per second (left axis) and average number of birds present (right axis, dashed grey) during the first 70 s of a mobbing event, averaged across sessions (n = 57). Lines show smoothed means for calls (magenta), calls given during a wing-lift (green), wing-lifts (blue), the proportion of birds actively signalling (dotted grey), and the proportion of birds being around the snake (present). **D**, odds ratios from a conditional logistic regression predicting active recruitment order, controlling for local baseline availability of each age–sex–rank class (n = 31 mobbing events, 104 individual joining events). Points represent odds ratios; horizontal lines denote 95% confidence intervals. The dashed vertical line indicates OR = 1 (probability equal to baseline availability). Dark grey, age–sex class comparisons; green, social rank comparison. Neither sex, age, nor rank predicted recruitment order. **E**, recruitment latency (response within 5 s, log scale) depending on the signal type that preceded a bird’s response (n = 1255 observations, 43 responders, 29 mobbing events). Coloured points show model-predicted means with 95% confidence intervals; grey points and connecting lines show individual-level means across trigger types. Birds responded significantly faster to a combined call–wing-lift display than to a call alone; latency following the onset of a wing-lift did also differ significantly from a call, but not from a combined call–wing-lift display. **F, left**, individual calling rate, and right, individual wing-lift proportion, both declining with group size. Model-predicted means (lines) with 95% confidence intervals (shaded bands) from generalized linear mixed models. Points show individual observations per session. Individual call rate declined significantly with group size; wing-lift proportion showed a negative trend. Call rate declined from 0.46 to 0.10 calls/s (77%), and wing-lift proportion from 0.23 to 0.10 (56%) across average group sizes 1–9. Squares, females; triangles, juveniles; diamonds, males. Significance indicators: *** p < 0.001, p < 0.01, * p < 0.05, • p < 0.1, ns p ≥ 0.05. Log-scaled axes are log-transformed.

## Methods

Details of the study site and population, as well as the experimental design, are described in Guggenberger et al., 2024.

### Study site and population

The study was conducted from February to August 2023 at the Shezaf Nature Reserve, located in the Arava region of the Negev Desert, a hyper-arid environment receiving approximately 30 mm of rainfall annually (Alaman et al., 2026; Anava et al., 2001; Keynan and Yosef, 2010). The Arabian babbler population at this site has been continuously monitored since 1971 and comprised approximately 120 individuals across 21 groups at the time of the study, with group sizes ranging from 3 to 18 birds. All individuals were marked with a unique colour ring combination to enable individual identification. Due to long-term habituation to human observers, individuals can be approached and observed at close range without inducing stress or altering behaviour (Zahavi, 1997).

### Experimental design

Snake-mobbing behaviour was elicited in ten groups of Arabian babblers, each comprising five to ten individuals. After confirming that group members were within visual or auditory range, a dummy viper (*Echis coloratus* or *Cerastes cerastes*) was placed partially embedded in the sand, and groups were recorded using a synchronized acoustic camera (SoundCam 2.0, CAE Software and Systems GmbH, Guetersloh, Germany) and a high-resolution RGB video camera (Canon EOS 5D) positioned approximately three meters from the model, enabling individual caller identification and postural analysis. Arabian babblers are known to hunt and kill other snake species, such as the Arabian cat snake (*Telescopus dhara*; Fig. S1). The dummy therefore had to appear convincing enough to elicit mobbing. Due to processing restrictions of the acoustic camera, mobbing was recorded in one-minute segments separated by ∼20-second pauses (session) to save the recording, and mobbing events ended when the group moved more than three meters from the model or stopped vocalizing for one minute.

#### Annotations

Acoustic camera recordings (Fig. 1A) of mobbing events were aligned with the RGB videos (Fig 1B) to 1min sections (sessions) due to technical constraints of the acoustic camera. The camera can only record for one minute, before it has to save the recordings in retrospect. Individual identity, age–sex class (adult female, adult male, juvenile), and dominance rank (dominant and subordinate) were known for all focal individuals. Juveniles were young birds till the age of six months. We only included recordings where at least one bird showed a wing-lift display and where all callers were identifiable (n=34 mobbing events). Three annotation pipelines were used to extract behavioural and acoustic data: Movement events were annotated and time-stamped using BORIS (Friard and Gamba, 2016). We marked when birds started and ended the wing-lift display. To determine caller identity by synchronizing the acoustic camera with RGB video recordings (25 fps), mobbing calls were annotated via Adobe Premiere Pro (Version 23.6.4). Call structure was analysed from audio recordings using Raven Pro 1.6.5 (K. Lisa Yang Center for Conservation Bioacoustics at the Cornell Lab of Ornithology, 2026). Calls were time-stamped, and acoustic parameters were extracted from the spectrograms. Movement videos (BORIS) [25fps], video-based call identity files (Adobe Premier Pro) [25fps], and acoustic (Raven Pro) annotation files in wave format [ms] were synchronized and aligned by time. Synchronization accuracy between calls in wave and video files (fps converted to ms) was 91.12%, and the average difference between call times in the two call files was 0.05 s (SD = 0.09 s). Calls that could not be aligned (8.88%) were excluded, resulting in 3199 identifiable calls.

### Analysis

All statistical analyses were conducted in R version 4.3.3 (R Core Team, 2024), using the packages glmmTMB (Brooks et al., 2025, 2017), lme4 (Bates et al., 2015), lmerTest (Kuznetsova et al., 2017), survival (Therneau, 2026), emmeans (Lenth, 2025), and DHARMa (Hartig, 2022). Figures were produced using ggplot2 (Wickham et al., 2025) and patchwork (Pedersen, 2025).

#### Recruitment

To determine how individual characteristics influenced the order in which birds entered the active mob, we used a conditional logistic regression model implemented as a discrete-choice risk-set design via clogit (survival). For each mobbing event, the initiator was defined as the first individual to perform any mobbing behaviour in the first session, and all remaining individuals were treated as followers, ranked by the timing of their first active participation across all sessions. Individuals were classified into five age-sex-rank classes (adult male dominant, adult male subordinate, adult female dominant, adult female subordinate, juvenile). At each recruitment step i, the risk set comprised all individuals not yet participating, with the chosen individual (arrival order = i) treated as the selected alternative. Models were grouped by recruitment step within each event, so that comparisons were made only among individuals available at the same point in the recruitment sequence, with the log-transformed proportion of each class in the risk set included as an offset to control for differential availability. Four pairwise contrasts of interest (adult female vs. adult male, juvenile vs. adult male, juvenile vs. adult female, adult subordinate vs. adult dominant) were estimated from this single model using simultaneous tests with single-step multiplicity adjustment, which corrects all four comparisons together to control the overall false-positive rate. Effects are reported as odds ratios (OR), representing the odds of a given individual being the next to join relative to others available in the risk set.

Recruitment latency was defined as the pause between an initiating signal (a call, a call during a wing-lift, or the onset of a wing-lift display) and the next signal produced by a different group member. We restricted responses to those occurring within 5 s of the initiating signal, since distance to the signal, and thus arrival time, could not be controlled for in the field, and slower responses may simply reflect birds still arriving rather than a true difference in latency. To examine how initiating signal type and responder age/sex class predicted recruitment latency, we fitted a generalized linear mixed model (glmmTMB) with a Gamma distribution (log link). The model included signal type and age/sex class as fixed effects, with responder identity, initiating individual identity, and mobbing event as random intercepts. Post-hoc pairwise comparisons were conducted using estimated marginal means (emmeans) with Tukey adjustments for multiple testing, and predicted latencies were back-transformed to the response scale (seconds). Model fit was assessed using DHARMa simulated residuals, with dispersion and outlier tests showing no significant violations.

#### Group size-rank- and age–sex class dependent signalling

We modelled call rate as the number of calls per individual per session, with session duration included as an offset to account for unequal observation windows; wing-lift proportion was standardized by dividing each individual’s cumulative wing-lift display duration by the total duration of their group’s session. To examine predictors of individual call rate and wing-lift proportion, we fitted two generalized linear mixed models (glmmTMB). Call rate was modelled with a negative binomial GLMM (log link), and wing-lift proportion was modelled with a beta distribution (logit link). Both models included dominance rank (dominant/subordinate), age-sex class (adult female/adult male/juvenile), group size, and time elapsed within the event as fixed effects. Group size was calculated as the mean number of birds present during the individual’s active behaviors. The elapsed time variable was modelled in raw seconds for the call rate model, but z-score standardized for the wing-lift model. Individual identity and event identity were included as random intercepts. Marginal predictions across the observed range of group sizes were obtained using emmeans, averaging over rank and age-sex class; call rate predictions were expressed as calls per second. Model fit was assessed using DHARMa simulated residuals, with dispersion and outlier tests showing no significant violations for either model.

#### Interplay and structure of acoustic and visual displays

Each wing-lift was classified as vocal or silent based on whether the individual producing the wing-lift also vocalized during that phase. The probability of a wing-lift being silent was modelled using a binomial GLMM (lme4), with age-sex class, dominance rank, group size and time elapsed, as fixed effects and individual identity and event identity as random intercepts. Wing-lift duration across successive displays within an event was modelled using a Gamma GLMM (glmmTMB, log link) with wing-lift order as a fixed effect and nested random intercepts for individuals within the event.

Inter-call pause duration was calculated as the time between consecutive calls from the same individual within the same session. Pauses longer than 5 s were excluded. To compare call structure between wing-lift and non-wing-lift contexts, we analysed peak frequency, call duration, and inter-call pause duration. Only individuals producing calls in both contexts were included (26 individuals, 35 sessions, 1871 calls). Call duration was log-transformed prior to analysis. Peak frequency and log-transformed call duration were modelled using linear mixed models (lme4 and lmerTest), and inter-call pause duration using a lognormal GLMM (glmmTMB). All three models included calling context (within vs. outside wing-lift) and sex/age class as fixed effects, with individual identity and session as random intercepts.

To examine how call structure changed across and within wing-lift displays, we fitted analogous models restricted to calls produced during wing-lifts (672 calls, 27 individuals, 33 sessions). Two sets of models were run for each acoustic variable: one with elapsed time since the individual’s first wing-lift in the mobbing event as the predictor (capturing change across the mobbing bout), and one with the call’s relative position within the wing-lift (0–1, from first to last call within a phase) as the predictor (capturing within-display change). Peak frequency and log-transformed call duration were modelled using linear mixed models (lme4 and lmerTest); inter-call pause duration was modelled using a Gamma GLMM (glmmTMB, log link). The peak frequency model additionally included wing-lift order as a covariate. All models included individual identity and session as random intercepts. Model-predicted means and curves with 95% confidence intervals were obtained using emmeans or computed directly from the fixed-effect design matrix and model variance-covariance structure. All models were assessed using DHARMa simulated residuals; dispersion and outlier tests indicated no significant violations.

## Results

### Recruitment

In total, we recorded 34 mobbing events across 9 groups in which birds showed visual displays. Birds readily joined the individual that detected the snake with 2.4 ± 1.7 birds joining the mob during the first minute (Mean±SD) (Fig. 1C). This number increased through recruitment to a maximum of 4.8 ± 1.6 individuals, with a mean of 3.7 ± 1.6 birds present throughout the mobbing events which lasted on average 2.4 ± 2 minutes. Of those, 1.1 ± 0.3 individuals were actively engaged in mobbing signalling at any given time. On average, all birds together emitted 1.03 ± 0.7 calls/s and performed 0.37 ± 0.36 wing-lift displays/s during mobbing. Juveniles called at a higher rate than adult females (0.32 vs. 0.19 calls/s; *z* = 2.08, *p* = 0.038; Tab.S3), while rank and sex of adult birds did not predict call rate or the proportion of wing-lift duration (all *p* > 0.14; Tab.S3).

Neither sex, rank, nor age predicted recruitment order of actively mobbing birds (adult females vs. adult males: p = 0.99; juveniles vs. adult males: p = 0.41; juveniles vs. adult females: OR = 0.47, p = 0.20; subordinates vs. dominants: p = 0.89; Fig. 1D; Tab. S1)

Recruitment latency while mobbing (defined as a response within 5 s to exclude birds still arriving) depended on signal type. Based on Tukey-adjusted post-hoc comparisons, birds responded with mobbing-related behaviour significantly faster to a combined call-wing-lift display than to a call alone (Gamma GLMM; z = -4.61, p < 0.001; predicted latency 0.61 s vs. 0.84 s; Fig. 1E; Tab. S2). Responses to a wing-lift display alone were also significantly faster than to a call alone (z = -2.76, p = 0.016). There was no significant difference in latency when comparing the combined display directly to the wing-lift display alone (z = -0.15, p = 0.988; Fig. 1E). The type of response also differed by trigger: a call alone (n = 868) predominantly recruited a further call (84.8% of responses), whereas both the combined call-wing-lift display (n = 352) and the wing-lift display alone (n = 75) predominantly recruited a combined call-wing-lift response (58.8% and 62.7%, respectively). Neither age nor sex predicted recruitment latency (all adjusted p ≥ 0.96; Tab. S2).

### Group size-dependent mobbing behaviour

Both individual calling and wing-lifting decreased as group size increased, independent of time elapsed (p > 0.6). Individual call rate declined significantly with each additional bird (negative binomial GLMM; z = −3.68, p < 0.001; Fig. 1F, left; Tab.S3), and wing-lift proportion (i.e., the proportion of time spent in this posture) showed a negative trend (beta GLMM; z = −1.74, p = 0.082; Fig. 1F, right; Tab.S3).

### The interplay between visual and acoustic signalling

Of 3199 calls recorded, 64.7% were produced without a simultaneous wing-lift display, while 35.3% occurred throughout the wing-lift. These calls were distributed broadly across the display but with notably fewer calls toward the display’s offset (KS test: D = 0.08, p < 0.001, Fig.S2). On the other hand, 80.2% of the 222 wing-lifts were performed with an accompanying call. The probability of performing a silent wing-lift was significantly higher in adult females (17%, z = 2.03, *p* = 0.042) and juveniles (25%, z = 2.26, p = 0.024) compared to adult males (4%, Fig. 2A, Tab. S3).

**Fig. 2.**
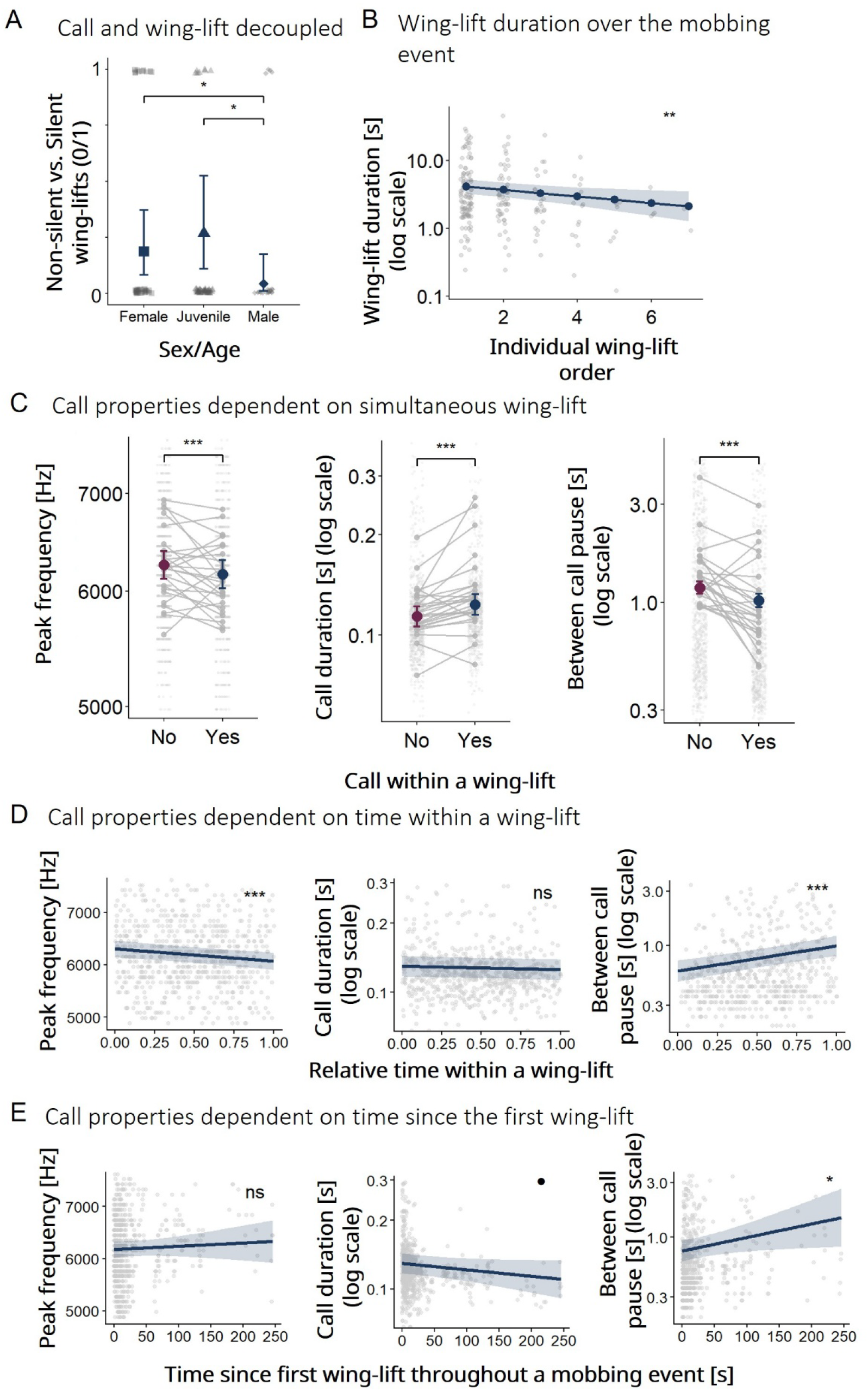
Interplay between mobbing acoustic and visual displays. **A**, probability of performing a silent wing-lift for each sex-age class. Adult females and juveniles were more likely to perform silent wing-lifts than adult males. Filled symbols show model-estimated means with 95% confidence intervals; grey points show raw observations, jittered horizontally. Squares, adult females; diamonds, adult males; triangles, juveniles, n = 222 wing-lifts, 39 individuals, 34 mobbing events. **B**, wing-lift duration as a function of successive display order within a mobbing session (n = 222 wing-lifts). Wing-lift duration decreased significantly with display order. Line shows model-predicted mean with 95% confidence interval (shaded); grey points show raw observations, jittered horizontally. **C**, Peak frequency, call duration, and between-call pause for calls produced outside (purple; n = 2,071) and within (blue; n = 1,128) wing-lift displays (26 individuals, 35 sessions). Wing-lift calls were lower in peak frequency, longer in duration, and preceded by shorter inter-call pauses. Filled circles show model-estimated means with 95% confidence intervals; grey points show raw observations; grey lines connect per-individual means. **D**, Acoustic properties as a function of the relative temporal position of a call within a wing-lift display (0 = start, 1 = end). Peak frequency declined toward the end of a display and inter-call pauses lengthened significantly; call duration was unaffected. **E**, Acoustic properties of wing-lift calls as a function of time elapsed since the first wing-lift in a mobbing event (n = 672 calls, 27 individuals, 33 events). Call duration showed a non-significant negative trend over time and peak frequency was unaffected, whereas inter-call pause duration lengthened significantly as the event progressed. In **D** and **E**, lines show model-predicted means with 95% confidence intervals (shaded); grey points show raw observations, jittered horizontally. Significance indicators: *** p < 0.001, ** p < 0.01, * p < 0.05, • p < 0.1, ns p ≥ 0.05. Log-scaled axes are log-transformed.

Calls produced during wing-lifts differed acoustically from calls produced without a wing-lift. Wing-lift calls frequency dropped from a mean of ∼6300 Hz to ∼6210 Hz, representing a significant ∼90 Hz decrease in peak frequency (LMM; t = −3.38, p < 0.001; Fig. 2C, left, Tab. S5). Additionally, these calls were ∼9.2 ms longer in duration (t = 6.46, p < 0.001; Fig. 2C, middle, Tab. S5) and were preceded by 0.17s shorter inter-call pauses (lognormal GLMM; z = −5.25, p < 0.001; Fig. 2C, right, Tab. S5).

Sex and age did not predict any of these acoustic properties (all p > 0.05, Tab. S5), indicating that babblers might not have a sexual dimorphism in voice, or at least not within the few properties we focussed on.

While the average frequency decrease was ∼90 Hz, this shift was also modulated by the call’s relative position within the wing-lift, with peak frequency declining by ∼241 Hz from display onset to offset – the shift was much larger towards the end of the wing lift (LMM; t = −3.49, p < 0.001; Fig. 2D, left, Tab. S6). Call position within a winglift did not predict call duration (p = 0.1; Fig. 2D, middle, Tab. S6), but inter-call pauses significantly lengthened by ∼0.40 s towards the end of a display (Gamma GLMM; z = 5.19, p < 0.001; Fig. 2D, right, Tab. S6).

The duration of each wing-lift decreased significantly with each successive display by the same individual (Gamma GLMM; z = −2.64, p = 0.008; Fig. 2B, Tab. S4). Call duration showed a negative non-significant trend over the course of a mobbing event (t = −1.89, p = 0.07; Fig. 2E, middle, Tab. S7), while inter-call pauses lengthened significantly with elapsed time (Gamma GLMM; z = 2.32, p = 0.02; Fig. 2E, right, Tab. S7). Peak frequency was unaffected by time (p = 0.45; Fig. 2E, left, Tab. S7).

## Discussion

Arabian babblers join a snake mob following the zwick alarm acoustic signals broadcasted by the individual who initiated the behaviour, usually the one who discovers the snake. Birds’ order of joining the mob within the same group was generally independent of rank, sex, or age. Similarly, in jackdaws (*Corvus monedula*), the sex of a calling colony member does not affect whether other members join the mobbing, unless the caller is an unfamiliar bird, in which case jackdaws are less likely to join female than male strangers (Woods et al., 2018).

Individual signalling efforts declined with group size (call rate decreased significantly and wing-lift proportion decreased marginally). During threatening events, familiar conspecifics can down-regulate an individual’s physiological stress response, a phenomenon known as social buffering. This effect depends on positive social bonds; in gregarious species, larger groups offer stronger protection than single companions (Hennessy et al., 2009). As more individuals join the mobbing, each bird’s perceived threat level may have decreased, lowering the internal drive to invest in defensive signalling. This physiological mechanism complements game-theoretical predictions of collective defense: as group size increases and personal predation risk is diluted, the cost of active participation outweighs the benefit. This lets individuals conserve energy by relying on the rest of the group to mob instead (Dugatkin and Godin, 1992; Flasskamp, 1994).

Our analysis suggests an asymmetric two-signal, two-receiver signalling system whereby conspecifics receive and integrate both signals, but the snake is limited to the visual one. Calls likely function as alarm-recruitment signals directed at conspecifics. The acoustic signal is ineffective against the snake itself: snakes lack an external ear and a tympanic middle ear, so they can only hear below 1000 Hz and are effectively deaf to the babblers’ ∼6200 Hz calls (Christensen et al., 2012). Instead, ambush snakes—like the Saharan sand viper (*Cerastes cerastes*)—rely on visual motion and ground-borne vibrations to localize prey, allowing them to hunt successfully even without chemosensory input (Young and Morain, 2002). Therefore, much like banner-tailed kangaroo rats (*Dipodomys spectabilis*) and California ground squirrels (*Otospermophilus beecheyi*), which utilize seismic foot-drumming and close-range visual tail-flagging to signal to ambush snakes, Arabian babblers likely utilize their visual wing-lift postures, and potentially the ground-borne vibrations generated by their constant movement, as reliable channels for signalling to these reptilian predators (Brownell, 1977; Mortimer, 2017; Randall and Matocq, 1997; Rundus et al., 2007). A babbler mob likely creates stress and a confusion effect for the snake — with multiple birds moving simultaneously around it, the snake’s ability to target and accurately strike any single individual is likely reduced (Ioannou et al., 2008; Olson et al., 2013). Babblers are group-living and have been observed directly attacking snakes collectively, including pecking at the tail (e.g., Arabian cat snake (*Telescopus dhara*), Fig. S1); our observations raise the possibility that the wing-lift display facilitates such attacks by distracting the snake while another individual strikes, though this remains untested. In our experiment, the dummy snake did not move and therefore probably shortened the mobbing effort. Babblers did not attack the coiled-up dummy viper, but eventually flew off. This indicates that vipers are either still seen as too dangerous and impossible to fend off, or that the dummy eventually became too weak a stimulus.

While most of the time these acoustic and visual signals are coupled, our dataset reveals an interesting context-dependent decoupling. Roughly 20% of the wing-lifts in our study were performed silently. This relative silencing may serve additional survival functions, as Zahavi, 1997 suggested; focussing on only the visual display could better prepare the bird for a sudden attack or escape. Visual-acoustic decoupling was modulated by both age and sex. Juveniles vocalized more overall, but performed 25% of their wing-lifts silently, a pattern likely reflecting inexperience as they refined their threat assessment (Guggenberger et al., 2024; Maklakov, 2002). In contrast, while silent wing-lifts are generally rare, adult females showed them more frequently than adult males (17% vs. 4%). Since females spend more time in snake-directed mobbing overall (Maklakov, 2002), dropping the call may imply that they are shifting effort from recruiting others to actively deterring and potentially attacking the predator.

Recruitment involves both physically joining a mob and stimulating others to signal. Our response time analysis measured the latency for a receiver to emit its own signal within a 5 s window, capturing this latter form of recruitment. Recruitment latency did not differ between wing-lift onset and the multimodal signal. The visual-only signal latency had a smaller sample (n = 75) than calls (n = 868) or multimodal signals (n = 352). Still, the results suggest the visual and multimodal displays act as effective triggers during close-range mobbing, indicating babblers process the visual stimulus alongside the acoustic one. A similar pattern of multimodal enhancement was documented in eastern grey squirrels (*Sciurus carolinensis*), where combined visual and acoustic alarm signals elicit stronger responses than either modality alone (Partan et al., 2009). In California ground squirrels (*Otospermophilus beecheyi*), tail flagging during snake encounters serves to attract nearby group members (Hersek and Owings, 1993). Ambush snakes in the sand are well camouflaged and easily overlooked, making visual displays an effective way to pinpoint the predator’s location for naive group members (Vail et al., 2013). Alternatively, since a bird’s call is physically altered during the motion of a wing-lift, the faster response towards the multimodal signal might be driven by the inherently different acoustic nature of these specific calls rather than the visual addition.

Arabian babbler “zwick” calls produced during wing-lifts differed acoustically from those produced without the visual display. During wing-lifts, individuals emitted significantly lower-frequency, elongated calls and emitted at faster rates, an acoustic shift that remained consistent regardless of sex or age. Birds can detect frequency changes of 1% or less (Dooling et al., 2000; Nagel et al., 2011). The 1.4–3.7% frequency drops we observed during wing-lifts cross that threshold, so birds could perceive them as distinct signals. The babblers’ slight 9.2 ms (8.54%) call lengthening falls below the standard 10% to 20% physiological threshold for avian duration discrimination, while the babblers’ 165 ms shortening in inter-call pause is much larger than the avian minimal gap-detection threshold (Dooling et al., 2000). This makes the increase in call rate and the decrease in frequency likely detectable to receivers, whereas the slight call lengthening might not be perceived, even though long outliers may be relevant.

Functionally, these dynamic acoustic changes are widely used across species to encode specific contextual information, such as the urgency of a threat or the caller’s motivation (Blumstein, 2025; Blumstein and Récapet, 2009; Sommer et al., 2012; Yorzinski and Vehrencamp, 2009). Lowering vocal frequency during high-threat encounters is often used functionally to exaggerate perceived body size or aggressive intent (de Boer et al., 2015). Domestic dogs (*Canis familiaris*) lower their growl frequency when facing a more threatening stranger, and orangutans (*P. pygmaeus wurmbii*) lower their alarm-call frequency in dangerous situations to sound larger than they are (Bálint et al., 2016; de Boer et al., 2015). Although snakes cannot detect “zwick” calls, raptors and other predators can, making it possible that the frequency drop may carry an evolutionary signature of predator deterrence while also encoding motivational information available to conspecifics during snake encounters (Zahavi, 1997). Focussing on snake mobbing and therefore on the acoustic signal for conspecific communication, signalling aggressive intent could function to coordinate the mob to harass and attack, showing the level of investment and risk taking, or both (Ioannou et al., 2008; Maklakov, 2002; Mine et al., 2022). Maklakov 2002 studied specifically how individual identity influences snake mobbing duration, also separating between reproductive stages, family-, and complex groups. As main dispersers, subordinates of family groups and females invested more time in mobbing, a pattern interpreted as self-advertisement (Maklakov, 2002). Simultaneously, accelerating the call rate is a well-documented strategy for encoding agitation and response urgency; for instance, the babblers’ 165 ms inter-call reduction even exceeds the interval decreases that American crows (*Corvus brachyrhynchos*) produce when signalling extreme danger during their escalation from stationary sitting to high-risk swooping attacks directly at a predator (Amit and Yovel, 2023; Yorzinski and Vehrencamp, 2009; Zhou et al., 2025).

Songbird vocal frequencies are influenced by the anatomical configuration of the upper vocal tract (Riede et al., 2006). The muscle contraction needed to hold the wing-lift posture likely alters these supralaryngeal structures, producing the drop in frequency and the timing shifts (Hoepfner and Goller, 2013; Riede et al., 2006). While birds accelerate calls during wing-lift, the inter-call pauses lengthen again towards the end of the display and also throughout the mobbing event. This lengthening likely reflects the accumulating physiological effort required to sustain the posture. For example, brown-headed cowbirds (*Molothrus ater*) introduce atypically long silent periods into their songs to compensate for the severe biomechanical and respiratory constraints caused by their own intense wing displays (Cooper and Goller, 2004; Hoepfner and Goller, 2013). Ultimately, playback experiments could shed a better light on the perception of calls produced during display vs. without.

## Conclusion

Snake-directed mobbing in Arabian babblers relies on an asymmetric two-signal, two-receiver system: the acoustic “zwick” call is inaudible to the snake and reaches only conspecifics, while the visual wing-lift display targets both. Babblers responded faster in close range mobbing to signals containing a visual component, multimodal or visual-only, than to calls alone. Recruitment order was independent of sex, age, and rank, while individual calling effort declined with group size, consistent with social buffering and predation-risk dilution. Wing-lifts were usually accompanied by calls, but silent wing-lifts occurred more often in juveniles and adult females, pointing to age- and sex-related variation in signal use. When coupled with wing-lifts, calls showed a lower peak frequency and faster call rate, changes that fall within known avian perceptual thresholds and are therefore likely detectable to receivers; this coupling was strongest for peak frequency toward the end of the display. Together, these results show that the two signal types are not independent channels but are dynamically linked, with the visual display shaping the acoustic one to produce a combined signal whose precise function — whether encoding motivation, coordinating the mob, or both — remains to be tested directly.

## Ethics statement

Institutional approval for this animal research was granted by Israeli Nature and Parks Authority (permit no. 2022/43151). All procedures complied with relevant institutional guidelines and local legislation.

## Data accessibility

Code and data will be made publicly available upon acceptance of the manuscript.

## Declaration of AI use

AI-assisted tools (Anthropic’s Claude and Google’s Gemini) were used for troubleshooting R code and for language editing in parts of the manuscript.

## Authors’ contributions

Marie Guggenberger: Writing – review & editing, Writing – original draft, Visualization, Software, Project administration, Methodology, Investigation, Formal analysis, Data curation, Conceptualization. Oded Keynan: Writing – review & editing, Supervision, Resources, Methodology, Conceptualization. Yossi Yovel: Writing – review & editing, Validation, Supervision, Resources, Methodology, Conceptualization.

## Acknowledgements

We acknowledge Nikola Dragić for providing photographic documentation of Arabian babblers preying on an Arabian cat snake (*Telescopus dhara*). Dorit Narisna and the Hazeva Field School provided accommodations during the fieldwork phase. Additionally, Yael Alon, Edgardo Santiago Rivera, Krista Oswald, and Alejandro Alaman provided practical and logistical support throughout and following the field research.

