## Supplemental Tables S1-7 and Figures S1+2 for "A two-signal, two-receiver signalling system: Snake mobbing in the Arabian babbler"

### Supplements

#### Figures

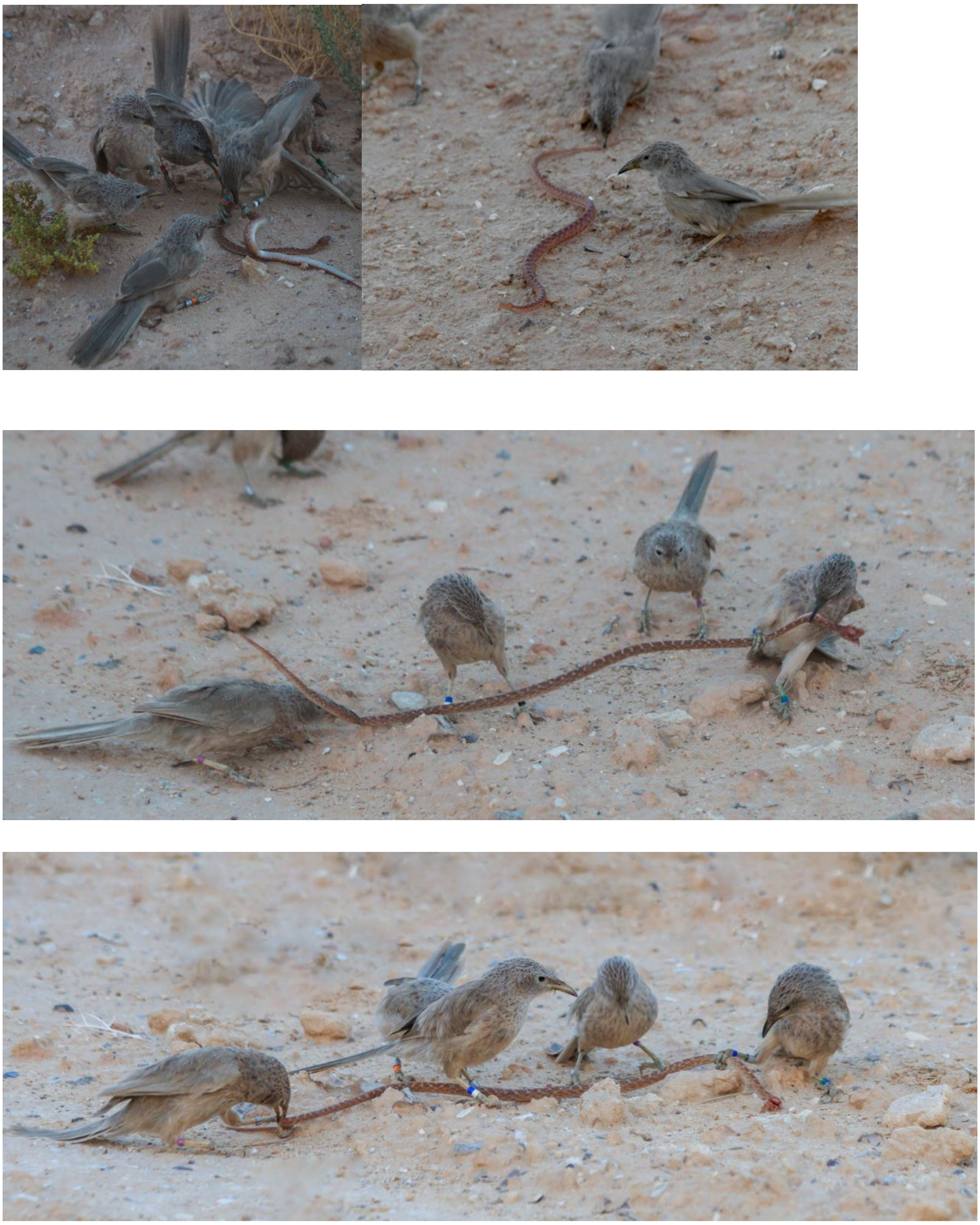

**Figure S1|** Arabian babblers preying on an Arabian cat snake (*Telescopus dhara*). Photo credit: Nikola Dragić.

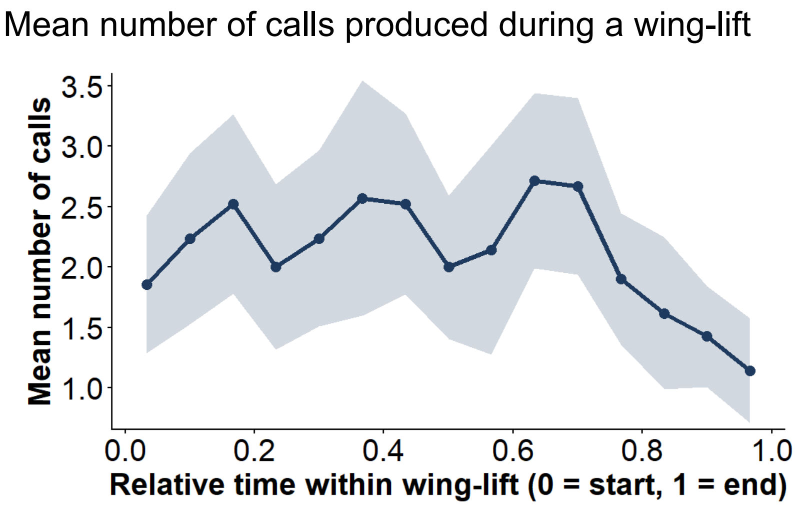

**Figure S2| Mean number of calls per wing-lift across relative time within the display** (n = 222 wing-lifts, 39 individuals, 15 bins). Calls were distributed broadly across the first ~75% of the display (mean range 1.95–2.67 calls per wing-lift) before declining markedly toward offset (bins 0.77–1.0: mean 1.19–1.86 calls). Points show binned means; shaded band shows ±1 SE.

#### Tables

**Table S1**: **Recruitment order dependent on identity** tested via conditional logistic regression.

| Predictor | Estimate | CI | Statistic | p-value |
|---|---|---|---|---|
| adult female vs. adult male | 1.13 | [0.45, 2.86] | 0.33 | 0.99 |
| juvenile vs. adult male | 0.53 | [0.18, 1.57] | -1.45 | 0.41 |
| juvenile vs. adult female | 0.47 | [0.17, 1.27] | -1.88 | 0.20 |
| adult subordinate vs. dominant | 1.31 | [0.48, 3.57] | 0.66 | 0.89 |

*p < 0.05; **p < 0.01; ***p < 0.001.

**Table S2**: **Recruitment latency dependent on multimodality of previous signal by another individual** tested using a generalized linear mixed model (GLMM, gamma distribution). Estimates and confidence intervals are on the log scale. Post-hoc pairwise comparisons include Tukey adjustments for multiple testing.

| Predictor | Estimate | CI | Statistic | p-value |
|---|---|---|---|---|
| call + wing-lift vs. call | -0.33 | [-0.47, -0.19] | -4.61 | <0.001*** |
| wing-lift onset vs. call | -0.35 | [-0.59, -0.10] | -2.76 | 0.02* |

| | | | | |
|---|---|---|---|---|
| juvenile vs. adult female | -0.01 | [-0.26, 0.24] | -0.11 | 0.99 |
| adult male vs. adult female | -0.03 | [-0.27, 0.20] | -0.27 | 0.96 |
| wing-lift onset vs. call + wing-lift | -0.02 | [-0.27, 0.23] | -0.15 | 0.99 |
| random effect: name (intercept) (variance) | 0.05 | | | |
| random effect: previous name (intercept) (variance) | 0.05 | | | |
| random effect: mobbing event (intercept) (variance) | 0.04 | | | |

*p < 0.05; **p < 0.01; ***p < 0.001.

**Table S3: Individual call rate, Individual wing-lift proportion, and probability of a silent wing-lift dependent on babbler identity** tested using generalized linear mixed models.

| Response variable | Predictor | Estimate | CI | Statistic | p-value |
|---|---|---|---|---|---|
| individual call rate | adult male vs. adult female | 1.37 | [0.87, 2.19] | 1.38 | 0.17 |
| individual call rate | group size | 0.83 | [0.75, 0.92] | -3.68 | < 0.001*** |
| individual call rate | time elapsed | 1 | [0.99,1.0] | -0.27 | 0.78 |
| individual call rate | juvenile vs. adult female | 1.72 | [1.05, 2.82] | 2.14 | 0.03* |
| individual call rate | subordinate vs. dominant | 1.38 | [0.88, 2.16] | 1.41 | 0.15 |
| individual call rate | random effect: mobbing event (intercept) (variance) | 0.05 | | | |
| individual call rate | random effect: name (intercept) (variance) | 0.25 | | | |
| individual wing-lift proportion | adult male vs. adult female | 0.81 | [0.52, 1.28] | -0.90 | 0.37 |
| individual wing-lift proportion | group size | 0.88 | [0.77, 1.02] | -1.74 | 0.08 |

| | | | | | |
|---|---|---|---|---|---|
| individual wing-lift proportion | time elapsed | 0.95 | [0.77, 1.18] | -0.44 | 0.66 |
| individual wing-lift proportion | juvenile vs. adult female | 1.23 | [0.79, 1.92] | 0.92 | 0.36 |
| individual wing-lift proportion | subordinate vs. dominant | 1.01 | [0.62, 1.63] | 0.03 | 0.98 |
| individual wing-lift proportion | random effect: mobbing event (intercept) (variance) | 0.07 | | | |
| individual wing-lift proportion | random effect: name (intercept) (variance) | 2.817e-10 | | | |
| probability of silent wing-lift | adult female vs. adult male | 4.94 | [1.06, 23.0] | 2.03 | 0.04* |
| probability of silent wing-lift | group size | 1.16 | [0.84, 1.62] | 0.89 | 0.37 |
| probability of silent wing-lift | time elapsed | 0.97 | [0.59, 1.59] | -0.13 | 0.89 |
| probability of silent wing-lift | juvenile vs. adult male | 7.89 | [1.31, 47.6] | 2.25 | 0.02* |
| probability of silent wing-lift | subordinate vs. dominant | 0.41 | [0.11, 1.64] | -1.26 | 0.21 |
| probability of silent wing-lift | random effect: mobbing event (intercept) (variance) | 0.75 | | | |
| probability of silent wing-lift | random effect: name (intercept) (variance) | 0.6 | | | |

*p < 0.05; **p < 0.01; ***p < 0.001.

**Table S4: Wing-lift duration dependent on order within the mobbing event** tested using a generalized linear mixed model (GLMM, gamma distribution).

| Predictor | Estimate | CI | Statistic | p-value |
|---|---|---|---|---|
| wing-lift order (successive display number) | 0.89 | [0.82, 0.97] | -2.64 | < 0.01** |
| random effect: name: mobbing event (intercept) (variance) | 0.18 | | | |

| | |
|---|---|
| random effect: mobbing event (intercept) (variance) | 0.3 |

*p < 0.05; **p < 0.01; ***p < 0.001.

**Table S5: Call properties dependent on wing-lift posture** tested using linear mixed models (LMM) for peak frequency and call duration; GLMM for pause duration.

| Response variable | Predictor | Estimate | CI | Statistic | p-value |
|---|---|---|---|---|---|
| call duration [s] | adult male vs. adult female | 0.02 | [0.0005, 0.04] | 1.98 | 0.06 |
| call duration [s] | juvenile vs. adult female | 0.002 | [-0.01, 0.02] | 0.21 | 0.83 |
| call duration [s] | wing-lift vs. normal | 0.01 | [0.006, 0.01] | 6.46 | < 0.01*** |
| call duration [s] | random effect: name (intercept) (variance) | 0.02 | | | |
| call duration [s] | random effect: session (intercept) (variance) | 0.01 | | | |
| pause [s] | adult male vs. adult female | -0.02 | [−0.16, 0.14] | -0.28 | 0.78 |
| pause [s] | juvenile vs. adult female | -0.07 | [−0.21, 0.10] | −0.83 | 0.41 |
| pause [s] | wing-lift vs. normal | -0.16 | [-0.22, -0.10] | -5.25 | < 0.01*** |
| pause [s] | random effect: name (intercept) (variance) | 0.01 | | | |
| pause [s] | random effect: session (intercept) (variance) | 0.01 | | | |
| peak frequency [Hz] | adult male vs. adult female | -149.79 | [-398.29, 100.55] | -1.16 | 0.26 |
| peak frequency [Hz] | juvenile vs. adult female | 18.3 | [-246.29, 284.93] | 0.13 | 0.90 |
| peak frequency [Hz] | wing-lift vs. normal | -88.62 | [-140.19, -37.49] | -3.38 | < 0.01*** |
| peak frequency [Hz] | random effect: name (intercept) (variance) | 64962.35 | | | |

| | | |
|---|---|---|
| peak frequency [Hz] | random effect: session (intercept) (variance) | 51447.4 7 |

*p < 0.05; **p < 0.01; ***p < 0.001.T-statistic for LMM, z for GLMM.

**Table S6: Call properties dependent on relative position within the wing-lift** tested using linear mixed models (LMM) for peak frequency and duration; GLMM for pause.

| Response variable | Predictor | Estimate | CI | Statistic | p-value |
|---|---|---|---|---|---|
| call duration [s] | relative position in wing-lift | −0.01 | [-0.01, 0.00] | −1.65 | 0.10 |
| call duration [s] | random effect: name (intercept) (variance) | 0.06 | | | |
| call duration [s] | random effect: session (intercept) (variance) | 0.01 | | | |
| pause [s] | relative position in wing-lift | 0.4 | [0.22, 0.60] | 5.19 | < 0.01*** |
| pause [s] | random effect: name (intercept) (variance) | 0.1 | | | |
| pause [s] | random effect: session (intercept) (variance) | 0.09 | | | |
| peak frequency [Hz] | relative position in wing-lift | −241.0 | [−376, −105] | −3.49 | < 0.01*** |
| peak frequency [Hz] | wing-lift order (period id) | 6.7 | [−54, 67.5] | 0.2 | 0.83 |
| peak frequency [Hz] | random effect: name (intercept) (variance) | 58405 | | | |
| peak frequency [Hz] | random effect: session (intercept) (variance) | 76022 | | | |

*p < 0.05; **p < 0.01; ***p < 0.001.

**Table S7: Call properties dependent on the bird's first wing-lift within the mobbing event** tested using linear mixed models (LMM) for peak frequency and duration; GLMM for pause.

| Response variable | Predictor | Estimate | CI | Statistic | p-value |
|---|---|---|---|---|---|

| | | | | | |
|---|---|---|---|---|---|
| call duration [s] | time since first wing-lift (s) | 0 | [-0.00, 0.00] | −1.89 | 0.07 |
| call duration [s] | random effect: name (intercept) (variance) | 0.06 | | | |
| call duration [s] | random effect: session (intercept) (variance) | 0.01 | | | |
| pause [s] | time since first wing-lift (s) | 0.002 | [0.0003, 0.004] | 2.32 | 0.02* |
| pause [s] | random effect: name (intercept) (variance) | 0.13 | | | |
| pause [s] | random effect: session (intercept) (variance) | 0.11 | | | |
| peak frequency [Hz] | time since first wing-lift (s) | 0.67 | [−1.1, 2.4] | 0.76 | 0.45 |
| peak frequency [Hz] | wing-lift order (period id) | 7.9 | [−59.1, 74.4] | 0.23 | 0.82 |
| peak frequency [Hz] | random effect: name (intercept) (variance) | 57093 | | | |
| peak frequency [Hz] | random effect: session (intercept) (variance) | 71269 | | | |

*p < 0.05; **p < 0.01; ***p < 0.001.
